# Selection of *Candida albicans* Trisomy during Oropharyngeal Infection Results in a Commensal-Like Phenotype

**DOI:** 10.1101/537340

**Authors:** Anja Forche, Norma V. Solis, Marc Swidergall, Robert Thomas, Alison Guyer, Annette Beach, Gareth A. Cromie, Giang T. Le, Emily Lowell, Norman Pavelka, Judith Berman, Aimeé M. Dudley, Anna Selmecki, Scott G. Filler

## Abstract

When the fungus *Candida albicans* proliferates in the oropharyngeal cavity during experimental oropharyngeal candidiasis (OPC), it undergoes large-scale genome changes at a much higher frequency than when it grows *in vitro.* Previously, we identified a specific whole chromosome amplification, trisomy of Chr 6 (Chr6x3), that was highly overrepresented among strains recovered from the tongues of mice with OPC. To determine the functional significance of this trisomy, we assessed the virulence of two Chr6 trisomic strains and a Chr5 trisomic strain in the mouse model of OPC. We also analyzed the expression of virulence-associated traits *in vitro.* All three trisomic strains exhibited characteristics of a commensal during OPC in mice. They achieved the same oral fungal burden as the diploid progenitor strain but caused significantly less weight loss and elicited a significantly lower inflammatory host response. *In vitro,* all three trisomic strains had reduced capacity to adhere to and invade oral epithelial cells and increased susceptibility to neutrophil killing. Whole genome sequencing of pre- and post-infection isolates found that the trisomies were usually maintained. Most post-infection isolates also contained *de novo* point mutations, but these were not conserved. While *in vitro* growth assays did not reveal phenotypes specific to *de novo* point mutations, they did reveal novel phenotypes specific to each lineage. These data reveal that during OPC, clones that are trisomic for Chr5 or Chr6 are selected and they facilitate a commensal-like phenotype.

## Introduction

Microbe-host interactions are highly complex. Following initial inoculation, multiple outcomes including colonization, commensalism, latency and disease are possible (Casadevall and Pirofski 2000; Casadevall 2017; Casadevall and Pirofski 2018). The immune status of the host is a key factor that determines the outcome of fungus-host interactions, especially for opportunistic fungi (Netea et al. 2015; Khannaa et al. 2016; Verma et al. 2017b). More recently, it has been appreciated that the genotype of the fungus also determines the outcome of this interaction. For example, in C. *albicans,* intra-species diversity among clinical strains results in differential modulation of fungus-host interactions (Schonherr et al. 2017). Similarly, intra-species diversity of *Cryptococcus neoformans* is associated with different clinical outcomes in patients with cryptococcal meningitis (Beale et al. 2015).

*C. albicans* is generally a clonal organism without conventional meiosis, and therefore the mechanisms to generate genotypic and phenotypic diversity are limited. Nonetheless, genomic analysis of a panel of clinical *C. albicans* strains revealed that many contained large-scale genome changes such as whole and segmental chromosome (Chr) aneuploidy and loss of heterozygosity (LOH), with frequent aneuploidy of Chrs 5 and 7. Interestingly, one strain with a loss-of-function mutation in *EGF1* exhibited decreased systemic virulence and increased gastrointestinal (GI) colonization in mouse models (Hirakawa et al. 2015). LOH events were commonly associated with acquisition of antifungal resistance, while aneuploidies appeared transiently in a study of oral strain series. In addition, several virulence-associated traits such as adherence and filamentation differed among the 43 strains tested (Ford et al. 2015).

Large-scale genome changes, including whole Chr and segmental LOH, also arise *in vitro* as a result of exposure to environmental stress such as nutrient limitation, oxidative stress, temperature and antifungal drug exposure (Rustchenko et al. 1994; Janbon et al. 1998; Gresham et al. 2008; Forche et al. 2011; Yona et al. 2012; Hill et al. 2013; Gerstein et al. 2015). Importantly, the frequency, type and extent of genomic changes is influenced by the nature and severity of the stressor (Forche et al. 2011).

Exposure to the host clearly represents the most complex stress that *C. albicans* encounters, and this interaction cannot be fully replicated *in vitro.* Previously, we determined that genotypic and phenotypic diversity appears as early as 1 day post-infection during both hematogenously disseminated and oropharyngeal candidiasis (OPC) in mouse models of infection (Forche et al. 2005; Forche et al. 2009; Forche et al. 2018). Genomic changes, in particular LOH, are 3 orders of magnitude more frequent *in vivo* compared to *in vitro* (Forche et al. 2009a; Ene et al. 2018; Forche et al. 2018). This strongly suggests that genome plasticity of the fungus may have a larger role in the fungus-host interaction than was previously appreciated. Our recent study of rapid C. *albicans* genome diversification during OPC identified a specific whole Chr amplification, trisomy of Chr 6 (Chr6x3), that was highly overrepresented among strains recovered from the tongues of mice after one round of infection. Chr6x3 was detected in 65% of mice and the allele combination ABB was 2-fold more common than the allele combination AAB (Forche et al. 2018).

Here, we tested the hypothesis that Chr6 trisomy is beneficial in the oral host environment and that strains with this genotype exhibit enhanced fitness compared to the progenitor during oropharyngeal infection. Strikingly, trisomy of Chr6x3 in two strains and trisomy of Chr5 (Chr5x3) in one strain all exhibited characteristics of commensals during oropharyngeal infection--they achieved the same oral fungal burden as the diploid progenitor strain yet caused significantly less weight loss, and they elicited a significantly lower inflammatory host response. In *vitro,* all three trisomic strains had reduced capacity to adhere to and invade oral epithelial cells. Whole genome sequencing showed that trisomies were mostly maintained, while point mutations that arose *de novo* in some lineages were unique to each lineage. Although *in vitro* growth assays did not reveal phenotypes specific to *de novo* point mutations, they did reveal novel phenotypes specific to each lineage under conditions that are relevant to fungus-host interactions and virulence potential. Taken together, our results reveal that Chr5x3 or Chr6x3 clones have a commensal-like phenotype that was apparently selected during OPC infection.

## Results

### Trisomic strains show a commensal phenotype in an oropharyngeal infection model

During oropharyngeal infection in mice, a specific trisomy, Chr6x3, was significantly enriched among strains and recovered from the majority of immunocompromised mice (Forche et al. 2018). The frequency of Chr6x3 increased over the course of infection (Fig. 1A) with the allele combination ABB occurring 2-fold more frequently than the AAB combination (Fig. 1B), suggesting that clones with trisomy of Chr6 have a general fitness advantage during OPC and that an extra copy of allele B may be more beneficial than an extra copy of allele A in this host niche. To test this hypothesis, we selected several strains that, based on whole genome karyotypes (produced using double digest restriction-site associated DNA sequencing (ddRADseq)), had acquired single trisomies as the only change compared to the diploid progenitor, strain YJB9318. Strains AF1275 and AF1485 both had acquired Chr6x3, the former with allele combination ABB (Chr6ABB) and the latter with allele combination AAB (Chr6AAB). Each strain was originally recovered from the oropharynx of the same mouse. Importantly, these strains had not been subjected to any selection regimes (e.g., *GAL1* counterselection-induced). A third strain, AF1773, that had acquired Chr5x3 (Chr5AAB) and a small LOH on Chr1 (due to selection for GAL1 LOH), served as control for a trisomy that did not involve Chr6. Of note, Chr5x3 was the second most common aneuploidy acquired in strains from the OPC model (Forche et al. 2018).

**Fig. 1.**
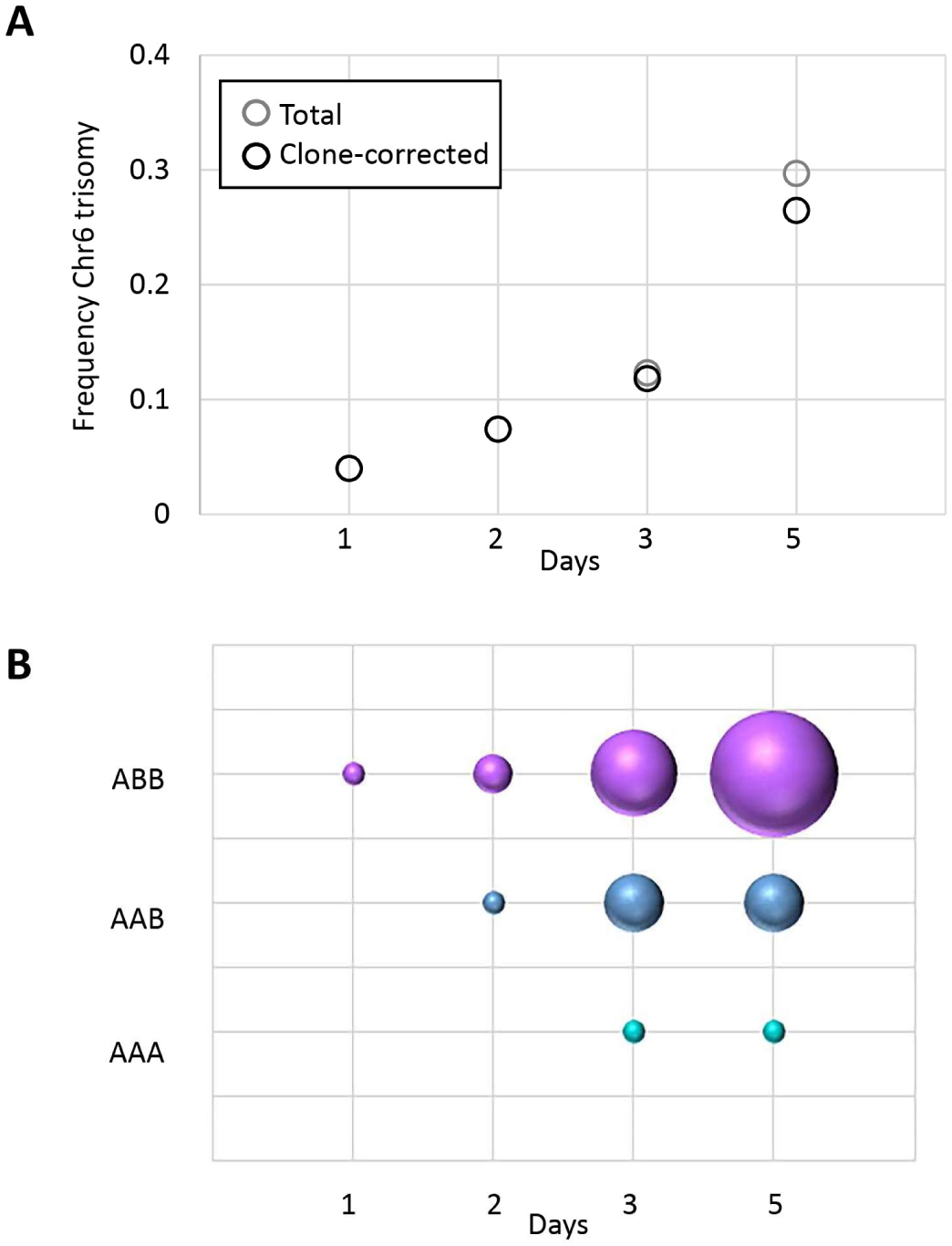
Chr6 trisomy ABB is overrepresented in isolates recovered from mice with OPC. (A) The frequency of Chr6 trisomy increases over the course of infection. (B) Among Chr6 trisomic strains, genotype ABB is the most frequent allele combination. For each genotype, symbol size is proportional to the frequency of isolation. Results are from the analysis of *C. albicans* colonies from 3-5 mice per time point as described in (Forche et al, 2018).

We assessed the virulence of the three trisomic strains (Chr6ABB, Chr6AAB, Chr5AAB) relative to their diploid progenitor during OPC in mice that had been immunosuppressed with cortisone acetate. Each strain was tested in 8 mice in two independent experiments for a total of 16 mice per strain. After 5 days of infection, the oral fungal burden of mice infected with all trisomic strains was similar to that of mice infected with the progenitor in terms of both Log CFU per gram of tissue (Fig. 2A) and histopathology (data not shown). The inflammatory response induced by the different strains was assessed using the myeoloperoxidase (MPO) content as a marker for the accumulation of phagocytes (neutrophils and macrophages) in the oral tissues (Glasgow et al. 2007; DE et al. 2009). Strikingly, immunocompromised mice infected with the trisomic strains had significantly lower tissue MPO levels relative to mice infected with the progenitor strain (Fig. 2B). Furthermore, mice infected with the Chr6 trisomic strains lost significantly less weight than mice infected with the progenitor on days 4 and 5 post-infection, while mice infected with the Chr5 trisomic strain showed significantly less weight loss on day 4, but not day 5 post-infection (Fig. 2C). Thus, the trisomic strains were able to proliferate to wild-type levels in the oropharynx, yet induced less inflammation and caused less disease, suggesting they promoted a commensal-like association with the immunosuppressed host.

**Fig. 2.**
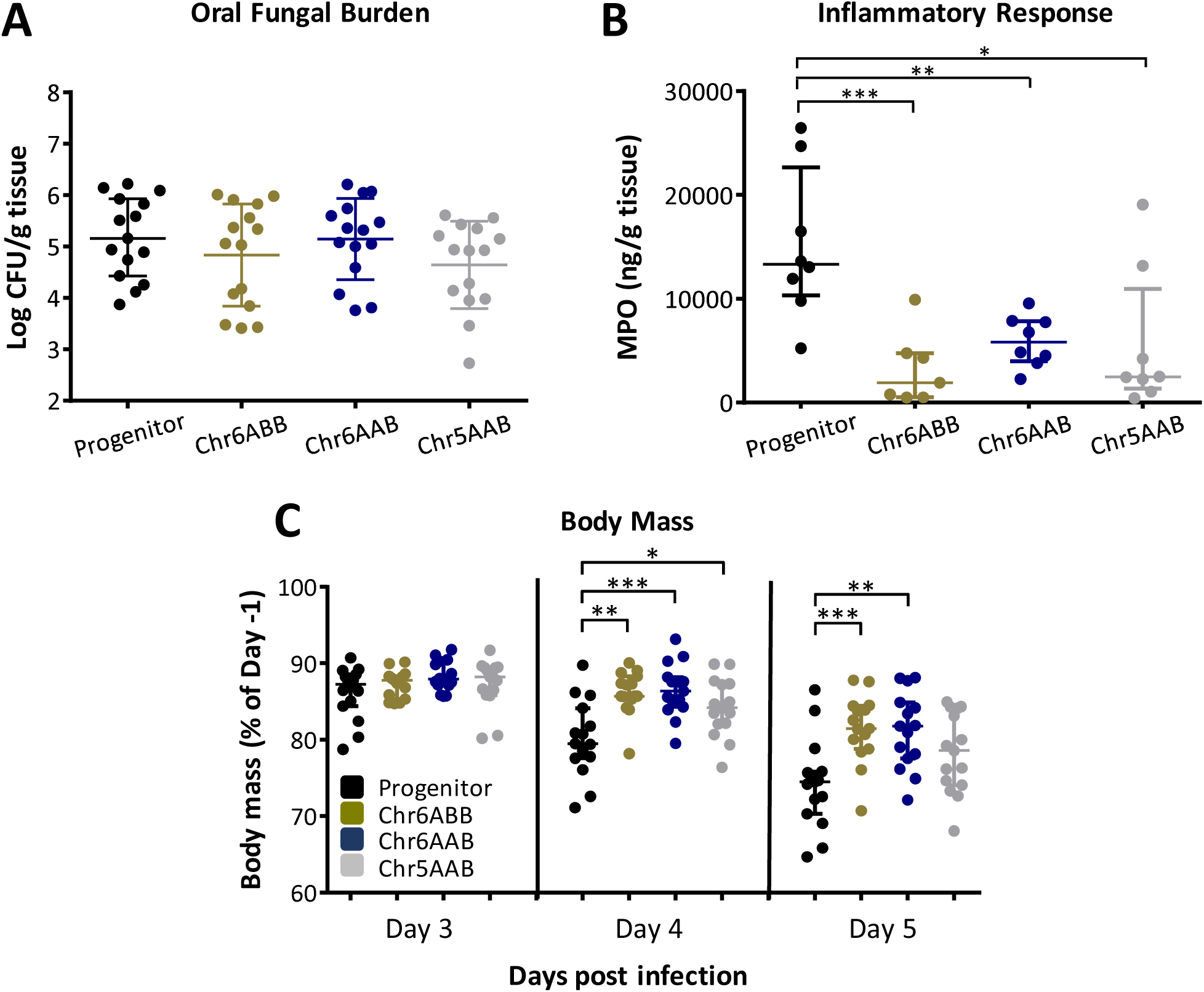
Trisomic strains exhibit commensal-like phenotype in immunosuppressed mice. (A) Oral fungal burden of mice after 5 d of infection with the indicated *C. albicans* strains. (B) Myeloperoxidase (MPO) levels in the oral tissues of mice after 3 d of infection with the indicated strains. (C) Body mass of mice after 3, 4 and 5 d of infection with the indicated strains. Results in (A) and (C) are the median interquartile range of combined data from 2 independent experiments, each using 8 mice per strain. Results in (B) are the median ± interquartile range of data from a single experiment with 8 mice per strain. *, p < 0.05, **, p < 0.01, ***, p < 0.001 by the Wilcoxon rank sum test.

### Chr6ABB behaves similar to the progenitor in an immunocompetent OPC model

Diverse clinical strains of C. *albicans* exhibit one of two phenotypes in the immunocompetent mouse model of OPC (Schonherr et al. 2017). The commensal-like group induces a relatively weak inflammatory response and persists in the oropharynx of immunocompetent mice for a prolonged time period. Others, such as blood isolate SC5314, induce a strong inflammatory response and are cleared from the oropharynx within 2-3 days. Of note, all strains used in our study were derived from strain SC5314. To ask if Chr6x3 induces a commensal-like phenotype in immunocompetent mice, we compared the Chr6ABB strain with the progenitor strain. The oral fungal burden and MPO content of mice infected with the Chr6ABB strain was similar to that of mice infected with the progenitor (Fig. 3A and B) after 1 day post-inoculation. By days 3 post-inoculation, the mice had cleared both strains of C. *albicans* from the oropharynx. Notably, the immunocompetent mice infected with the Chr6ABB strain lost significantly less weight than the mice infected with the progenitor (Fig. 3C), again indicating less severe disease associated with Chr6x3.

**Fig. 3.**
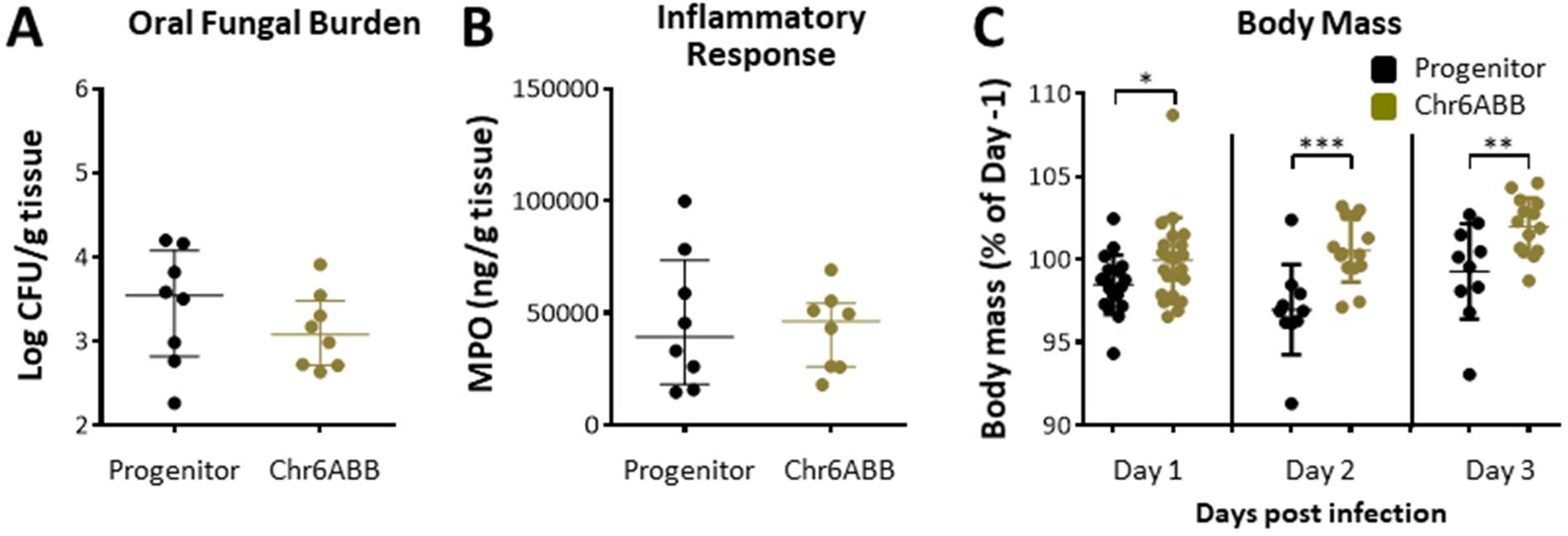
The Chr6ABB strain has a commensal-like phenotype in immunocompetent mice. (A) Oral fungal burden of immunocompetent mice after 1 d of infection with the progenitor and Chr6ABB strains. (B) MPO levels in the oral tissues of the mice after 1 d of infection with the indicated strains. (C) Body mass of the mice after 1, 2 and 3 d of infection with the indicated strains. Results in (A) and (B) are the median ± interquartile range of data from a single experiment, each using 8 mice per strain. Results in (C) are the median ± interquartile ranges of data from a single experiment with 18-22 mice per strain on d 1 and 10-14 mice per strain on d 2 and 3. *, p < 0.05, **, p < 0.01, ***, p < 0.001 by the Wilcoxon rank sum test.

### Infection with the Chr6ABB strain induced a weaker chemokine and IL-17A response

To further analyze the inflammatory response induced by the Chr6ABB strain, we determined the profile of chemokines, pro-inflammatory cytokines, and alarmins in the oral tissues of mice infected with this strain relative to the progenitor. In the immunocompetent mice, the Chr6ABB strain induced significantly less CCL3, CXCL1 (KC), IL-1β, IL-17A, and the p19 subunit of IL-23 in the oral tissues relative to the progenitor strain after 1 day of infection (Fig. 4A). The Chr6ABB strain also induced a weaker inflammatory response in corticosteroid-immunosuppressed mice after 5 days of infection (Fig. 4B). Collectively, these results indicate that the Chr6ABB strain induced an attenuated inflammatory response in both immunocompetent and immunosuppressed mice despite proliferating in the oropharynx to the same extent as the progenitor.

**Fig. 4.**
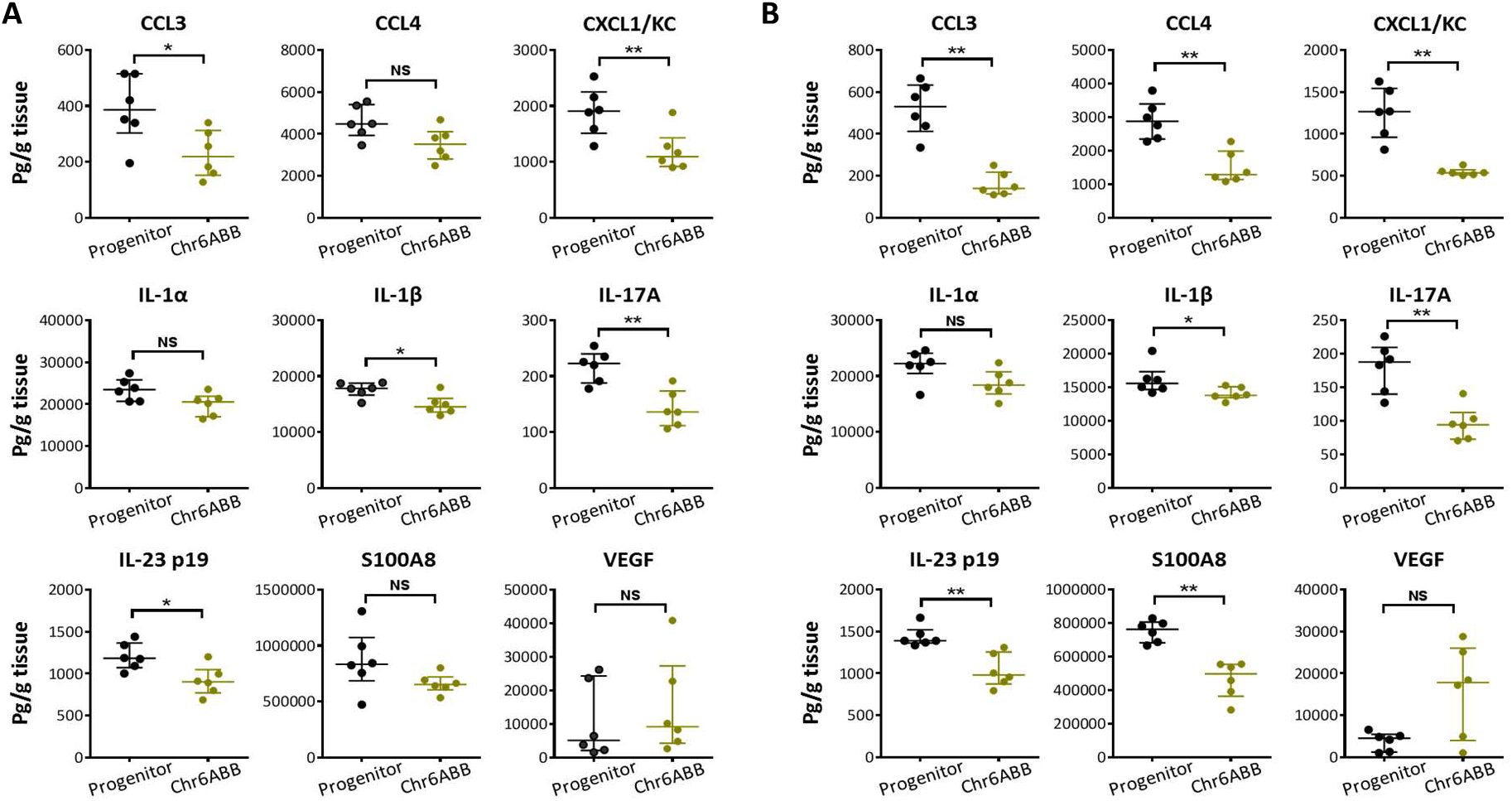
Oral infection with the Chr6ABB strain induced less production of inflammatory mediators in the oral tissues of immunocompetent mice (A) after 1 d of infection and immunosuppressed mice (B) after 5 d. Results are the median ± interquartile ranges of data from a single experiment with 6 mice per strain. *, p < 0.05, **, p < 0.01, ***, p < 0.001 NS, not significant, by the Wilcoxon rank sum test, corrected for multiple comparisons.

### The trisomic strains had reduced adherence to and invasion of oral epithelial cells, and were more susceptible to neutrophil killing

We tested the capacity of the trisomic strains to adhere to, invade, and damage oral epithelial cells, as well as their susceptibility to killing by human neutrophils, *in vitro.* The number of epithelial cell-associated organisms, a measure of adherence, was significantly reduced for the trisomic strains relative to the progenitor strain (Fig. 5A). The trisomic strains also were endocytosed poorly as compared to the progenitor strain (Fig. 5B). The two Chr6x3 strains (Chr6ABB and Chr6AAB) induced a similar extent of epithelial cell damage relative to the progenitor, while the Chr5x3 strain caused significantly less damage (Fig. 5C). Surprisingly, all three trisomic strains were more susceptible than the progenitor strain to neutrophil killing (Fig. 5D). Thus, strains with trisomy of Chr5 or Chr6 have reduced epithelial cell adherence and invasion as well as increased susceptibility to neutrophil killing *in vitro.*

**Fig. 5.**
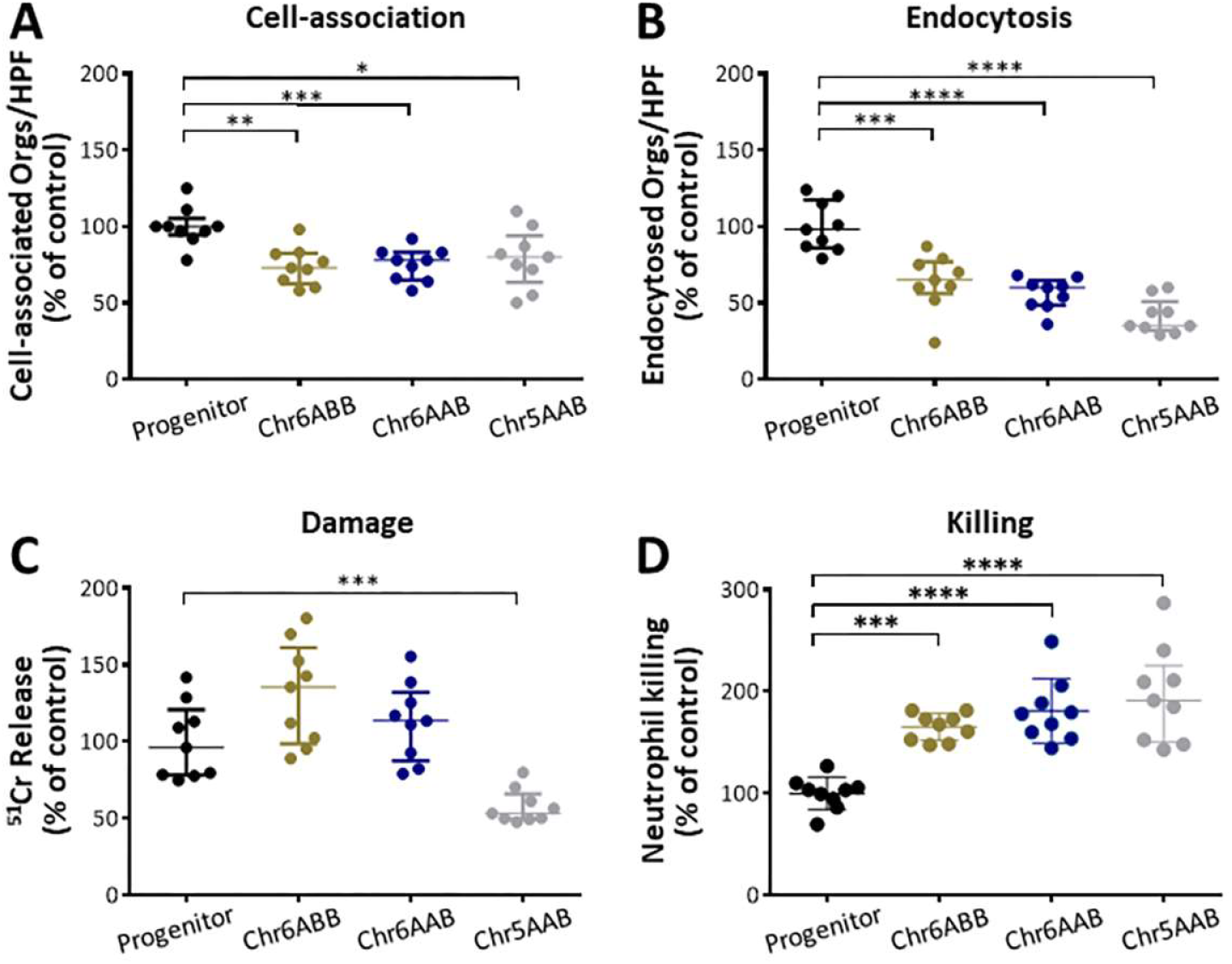
Effects of trisomy on the host cell interactions of *C. albicans In vitro*. (A) All trisomic strains exhibit significant defects in adherence to (A) and endocytosis by (B) oral epithelial cells. C. Only Chr5AAB shows significantly reduced damage of epithelial cells compared to progenitor D. Results are the median ± interquartile range of data from a single experiment with 6 mice per strain. *, p < 0.05, **, p < 0.01, ***, p < 0.001 ****, p < 0.0001, by the Wilcoxon rank sum test. HPF, high-power field; Orgs, organisms.

### Phenotypes common and unique to each trisomic lineage

We performed growth assays by spot dilution for the original trisomic strains (Forche et al. 2018) as well as for several single colony isolates recovered after oral infection. In addition to testing 17 different growth conditions at both 30°C and 37°C, we determined lipase/phospholipase hydrolytic activity on egg yolk agar (Fig.6, Fig.S1). Under the majority of conditions, the trisomic strains grew similarly to the progenitor strain. However, the Chr6x3 strains were less susceptible than the progenitor to SDS and rapamycin, and more susceptible to protamine sulfate (Fig. 6A, Fig.S1). By contrast, the Chr5AAB strain was more susceptible to rapamycin, but less susceptible to protamine sulfate. Interestingly, all three strains showed less filamentous growth that the progenitor (spots were smooth or only slightly wrinkled vs wrinkled progenitor colonies) under a variety of conditions (Fig. 6B, Fig.S1). These observations were strain-specific, with Chr5AAB showing the fewest defects and Chr6ABB the most defects in filamentous growth (Fig.6B, Fig.S1). The Chr6x3 strains also had reduced extracellular lipase/phospholipase hydrolytic activity relative to the progenitor strain (Fig. 6C). Importantly, strains AF1852 and AF1942, both of which lost one Chr6B allele during oral infection, exhibited progenitor phenotypes under all conditions tested (Fig.6, Fig.S1). This result strongly supports the idea that the extra copy of Chr6B was necessary and sufficient to induce the observed phenotypes in the Chr6x3 strain.

**Fig. 6.**
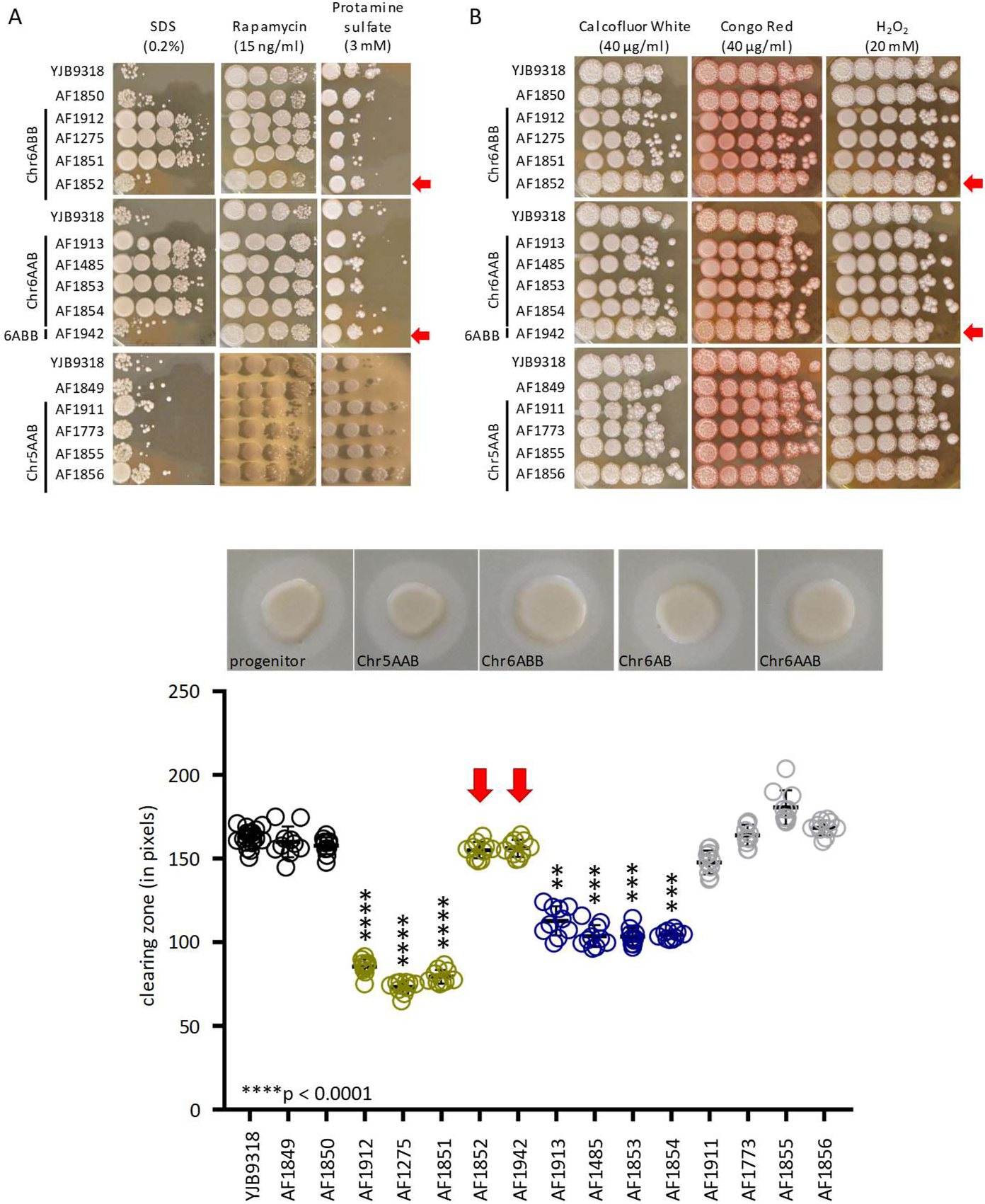
Phenotypic screening reveals lineage-specific trait changes. Note that AF1852 and AF1942, which lost one Chr6B allele, exhibit progenitor phenotypes under all conditions tested (see red arrows). A. Spot dilution assays at 37°C show differences in growth compared to the progenitor and distinct differences between Chr5 and Chr6 trisomic strains for three conditions. All images were taken on Day 3. See Fig.S1A for complete growth profiles. B. Both Chr6 lineages show defects in filamentous growth at 37°C specifically under cell wall and oxidative stress. Shown are 3 representative conditions. See Fig.S1 B for complete profiles. C. Both Chr6 lineages show significantly lower bulk extracellular lipase and phospholipase activity compared to progenitor. Top images are representative images showing clearing zone around cell spots.

### Trisomies in Chr6 were mostly maintained during OPC

Because strains with aneuploid chromosomes are often unstable, we measured strain ploidy levels both before and after OPC infection, using a qPCR approach that sampled 4 markers along each of the eight chromosomes from inoculum streaks, from single colonies of the inoculum, and from single colonies recovered from different mice after OPC. At the population level (streak), all 4 strains maintained their original karyotype over the three days of inoculum preparation (Fig. S2). However, when single colonies were analyzed, 1 of 5 colonies of the Chr6ABB strain had lost Chr6 trisomy, indicating that Chr6x3 is not very stable, and raising the possibility that the initial inoculum of this strain was a mixed population. Next, we analyzed multiple colonies from all four strains to ask if any large-scale genomic changes had occurred during infection. No novel aneuploidies were found in any of the analyzed strains (Fig.7A, Fig.S2). In total, 27% (6/22) of individuals from the Chr6ABB strain background had lost the trisomy during the course of the OPC infection experiment. Similarly, 31% (5/16) of the strains from the Chr6AAB strain background were no longer trisomic. By contrast, all post-infection strains of the progenitor and of the Chr5AAB strains had maintained their original karyotypes (see also below). Thus, Chr6x3 appears to be more unstable than Chr5AAB and thus likely requires positive selection for its maintenance in the OPC model.

**Fig. 7.**
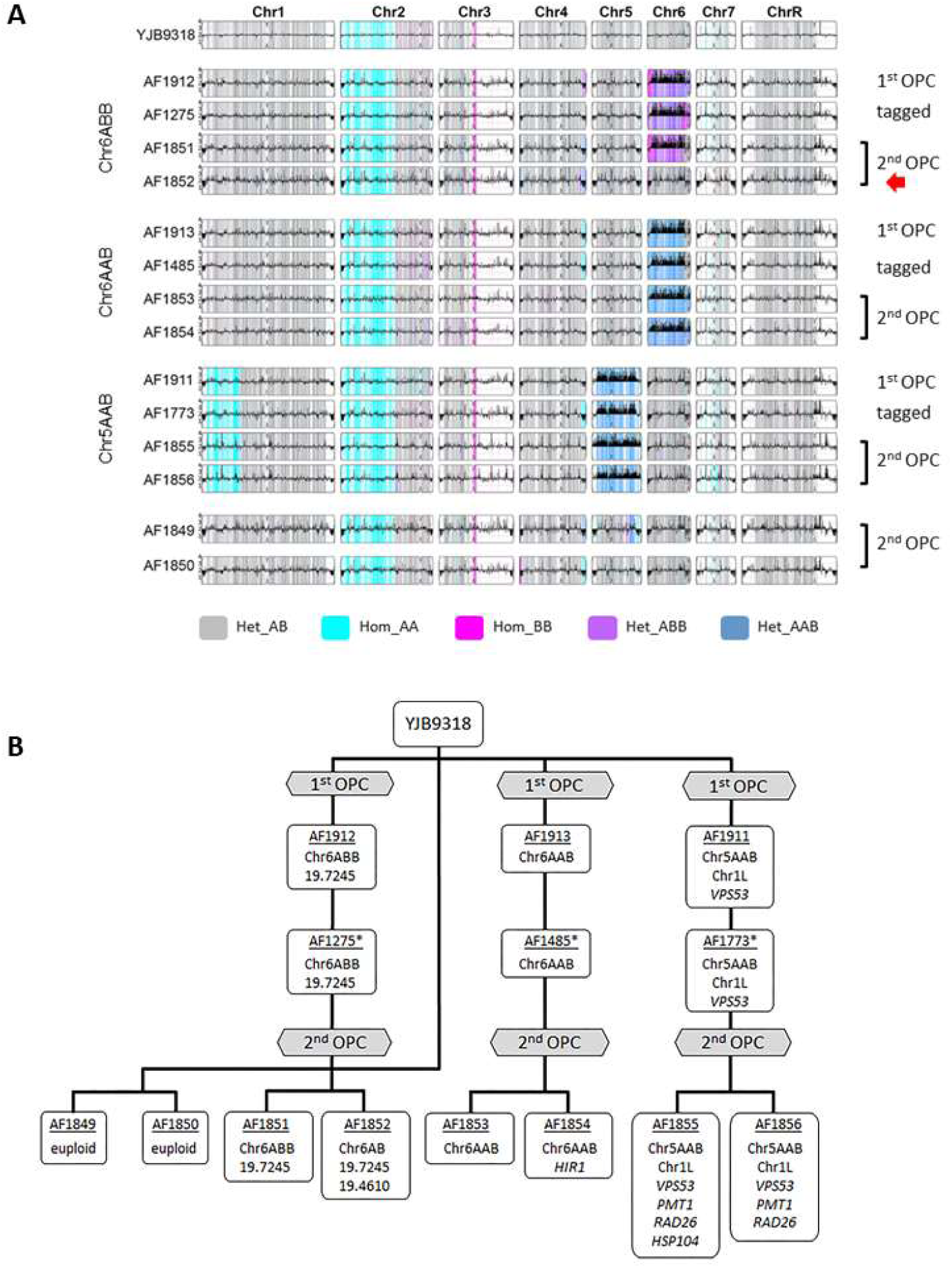
Whole genome sequence analysis of the trisomic strains. (A) The genomes of most strains were stably maintained during transformation and second oral infection. YMAPs showing the local copy number estimates for strains from which whole genome sequence data were obtained. Note that strain AF1852 had lost one copy of Chr6B. Het, heterozygous; hom, homozygous; 1^st^ OPC, original oral infection (Forche et al. 2018); 2^nd^ OPC, strains isolated from mice in the current experiments. (B) *De novo* point mutations arose during the 2^nd^ OPC but were not shared among trisomic lineages. Diagram of the strain lineages showing the genomic changes and the SNPs in the indicated genes (see Table S2 for detailed information on SNPs); * Tagged with unique barcode at the NEUT5L locus.

### Trisomic strains acquired relatively few *de novo* point mutations during OPC, and these were not conserved

SNPs, INDELS, LOH and CNVs were identified in the post-OPC isolates using Illumina whole genome sequencing (see Fig 7A). Maintenance and loss of aneuploid chromosomes, detected by qPCR, were confirmed for all strains (Fig. 7B). Importantly, the karyotypes of these strains were stable during transformation and during 5 days of OPC in all strains except for strain AF1852, which was a post-infection Chr6ABB strain (Fig. 7B). No *de novo* large-scale LOH events were observed. We identified a total of 8 de novo SNPs/Indels, all of which were heterozygous; we verified each SNP by Sanger sequencing of the parent and evolved strains (Fig. 7A, Table S2). Seven of these SNPs were located in coding regions and one was intergenic (Table S2). Thus, de novo mutations were identified in all three lineages, with none of them were recurrent (shared among strains). These results strongly suggest that the commensal-like phenotype was indeed due to whole Chr aneuploidies.

## Discussion

When C. *albicans* infects a mammalian host, it is exposed to a variety of different stressors that vary with both the anatomic niche and the duration of infection. Many stressors, such as reactive oxygen species produced by immune cells, antimicrobial peptides, and sequestration of micronutrients are generated by the host. During infection of non-sterile mucosal surfaces, the fungus must also withstand stressors generated by the competing bacterial microbiota. The magnitude of stress experienced by C. *albicans* at a specific infection site is likely dependent on whether the organism is growing as a commensal or a pathogen. For example, when C. *albicans* grows in the oropharynx as a commensal, it induces only a weak host response and is thus unlikely to experience significant oxidant stress induced by activated phagocytes. By contrast, when C. *albicans* overgrows and causes OPC, the massive influx of neutrophils subjects the organism to substantial oxidant stress (Swidergall and Filler 2017; Verma et al. 2017a). The multiplicity of stressors encountered by C. *albicans* necessitates that it adapts to specific host micro niches.

Aneuploidy and LOH of C. *albicans* generated *in vivo* is thought to enhance the survival and proliferation of the fungus within the host (Forche et al. 2005; Forche et al. 2009; Ene et al. 2018; Forche et al. 2018; Tso et al. 2018), a hypothesis that has been tested only in a mouse model of GI colonization (Ene et al. 2018; Tso et al. 2018). In particular, experimental evolution in the mouse GI tract selects for C. *albicans* variants harboring whole-chromosome as well as segmental aneuploidy and LOH events (Ene et al. 2018; Tso et al. 2018) and results in commensal, attenuated strains that protect their host against subsequent infection with a virulent C. *albicans* strain (Tso et al. 2018). The overrepresentation of strains with Chr6x3 in our earlier study (Forche et al. 2018), despite the instability of the trisomic state, suggests that this trisomy provides fungal cells with a competitive advantage during OPC. The results presented here demonstrate that trisomies of Chr6 and Chr5 enable *C. albicans* to infect the oropharynx at high levels, yet causes less disease. Specifically, when immunosuppressed mice were orally inoculated with the trisomic strains, their oral fungal burden was similar to that of mice infected with the diploid progenitor strain. At the same time, the trisomic strains induced a lower inflammatory response in terms of reduced phagocyte accumulation and decreased cytokine levels, which resulted in less weight loss than in mice infected with the progenitor strain. These results suggest that trisomy in either Chr6 or Chr5 causes *C. albicans* to begin to assume a commensal-like phenotype.

To further explore the mechanism of the commensal-like phenotype of the trisomic strains, we analyzed the host response to OPC caused by the Chr6 ABB strain relative to the progenitor strain. In both immunocompetent and immunosuppressed mice, the Chr6 ABB strain induced lower production of multiple chemokines and pro-inflammatory cytokines. The decreased levels of CCL3 and CXCL1, which are neutrophil chemoattractants, provides an explanation for the reduced levels of the phagocyte marker MPO in the oral tissues of mice infected with the trisomic strains. Reduced levels of IL-17A and IL-23, which play key roles in the host defense against OPC (Conti et al. 2009), also suggest that the trisomic strain induced less disease during OPC and are consistent with the commensal-like phenotype of the trisomic strains.

A potential explanation for the reduced pathogenicity of the Chr6x3 and Chr5x3 strains is provided by our *in vitro* studies: trisomic isolates had reduced capacity to adhere to and invade oral epithelial cells. C. *albicans* expresses a multitude of adhesins that mediate the binding of the fungus to epithelial cells (reviewed in (Höfs et al. 2016)). One of these adhesins is Als3, which also functions as an invasin (Phan et al. 2007; Zhu et al. 2012). One possible explanation for the reduced adherence and invasion of the trisomic strains is that they have reduced surface expression of Als3. However, by flow cytometric analysis of hyphae stained with an anti-Als3 antibody, we found that Als3 surface expression of all three trisomic strains was similar to that of the progenitor strain (Solis and Filler, unpublished data). These results suggest that the adherence and invasion defects of the trisomic strains are due to reduced expression of one or more adhesin(s) and invasin(s) other than Als3.

When Schonherr et *al.,* (Schonherr et al. 2017) investigated the epithelial cell interactions of different strains of *C. albicans*, they found that the ones with the commensal phenotype caused less damage to epithelial cells *in vitro* than did the pathogenic strains. We found that only the Chr5 AAB strain had reduced capacity to damage the epithelial cells; both Chr6 trisomic strains induced wild-type levels of epithelial cell damage. Thus, reduction of *in vitro* epithelial cell damage did not predict the commensal-like phenotype of the Chr6 trisomic strains.

Surprisingly, all three trisomic strains had increased susceptibility to neutrophil killing, yet the oral fungal burden of mice infected with these strains was similar to that of mice infected with the progenitor strain. We speculate that the trisomic strains were able persist in the oropharynx in high numbers because of the defect in phagocyte recruitment.

Whole genome sequence analysis did not identify any common point mutations among the trisomic lineages. This result supports the concept that the trisomic state of specific chromosomes, rather than specific point mutations, resulted in the commensal-like phenotype. In a recent study, trisomy of Chr7 (Chr7x3) was found to be common in *C. albicans* strains after *in vivo* passage in a mouse model of GI colonization (Ene et al. 2018). Chr7x3 arose in three different strain backgrounds and conferred a fitness (growth) advantage over the respective diploid progenitors in 1:1 *in vivo* competition experiments. Thus, the development of trisomy in specific chromosomes appears to influence the capacity of *C. albicans* to persist in distinct anatomic niches.

Chr6 harbors many genes important for filamentous growth, adhesion and hydrolytic enzyme production (Skrzypek et al. 2017). Importantly, strains that lost the extra copy of the Chr6B allele from the Chr6ABB strains reverted to the parental phenotypes. These data support the idea that the commensal phenotype is a multi-gene trait that is fostered by an extra copy of Chr6B. Allelic imbalance in the trisomic strains may also affect cell wall architecture and/or composition, which would then alter immune recognition by the host. Accordingly, the amount of exposed β-glucan on the surface of fungal cells was strongly predictive of competitive fitness in the mouse GI tract (Ene et al. 2018). Furthermore, fungal cell wall architecture, rather than cell wall composition, determines the ability of fungi to colonize the GI tract (Sem et al. 2016). Finally, a recent study on functional divergence of filamentous growth regulation in C. *albicans* found that Flo8 overexpression was sufficient to restore filamentation in a *mfg1/mfg1* mutant (Polvi et al. 2019). Importantly, Flo8 is located on Chr6 and *in vitro* evolution of three *mfg1/mfg1* lineages *in vitro* resulted in trisomy of Chr6 (Polvi et al. 2019).

Taken together, our data indicate that specific whole chromosome aneuploidies alter several related virulence-associated traits that affect how the host recognizes and responds to C. *albicans* during oropharyngeal infection, thereby inducing a commensal-like phenotype. Because both the *in vivo* (commensal) and *in vitro* phenotypes are likely the result of allelic imbalance of specific genes on the trisomic chromosomes, rather than due to whole chromosome trisomy, it will be imperative to identify those genes that, when present in an extra copy, enhance the capacity of *C. albicans* to interact with the host and survive in diverse anatomic sites.

## MATERIALS AND METHODS

### Strains used in this study

Strains are listed in Table S2 and were stored at −80°C in 50% glycerol.

### Ethics statement

All animal work was approved by the Institutional Animal Care and Use Committee (IACUC) of the Los Angeles Biomedical Research Institute. The collection of blood from human volunteers for neutrophil isolation was also approved by the Institutional Review Board of the Los Angeles Biomedical Research Institute.

### Mouse model of OPC

The pathogenicity of the C. *albicans* strains during OPC was determined in both immunocompromised and immunocompetent male Balb/c mice following our standard protocol (Solis and Filler 2012). When immunocompromised mice were used, cortisone acetate (2.25 mg/kg) was administered subcutaneously on days −1, 1, and 3 (Solis and Filler 2012). For inoculation, the animals were sedated with ketamine and xylazine, and a swab saturated with 10^6^ C. *albicans* cells was placed sublingually for 75 min. Immunocompetent mice were inoculated similarly, except that the swab was saturated with 2 × 10^7^ organisms (Conti et al. 2016; Solis et al. 2017). The immunocompromised and immunocompetent mice were sacrificed at 3 and 5 d and 1 and 3 d post-infection, respectively. After sacrifice, the tongue and attached tissues were harvested and divided longitudinally. One hemisection was weighed, homogenized, and quantitatively cultured. The other was embedded in paraffin, after which thin sections were prepared and then stained with periodic acid-Schiff stain (PAS).

### Human cell line

The human oral epithelial cell line OKF6/TERT-2 was kindly provided by J. Rheinwald (Harvard University, Cambridge, MA) (Dickson et al. 2000) and was cultured as previously described (Phan et al. 2007).

### Host cell damage assay

The extent of oral epithelial cell damage caused by different C. *albicans* strains was determined using our previously described ^51^Cr release assay (Solis et al. 2017). Briefly, OKF6/TERT-2 cells were grown to 95% confluence in 96-well tissue culture plates with detachable wells (Corning) and loaded with 5 µCi/ml Na_2_^51^CrO_4_ (PerkinElmer) overnight. After rinsing the cells to remove the unincorporated ^51^Cr by rinsing, they were infected with 6 × 10^5^ C. *albicans* cells. After 7 h, the amount of ^51^Cr released into the medium and retained by the cells was determined by gamma counting. Each experiment was performed three times in triplicate.

### Measurement of *C. albicans* epithelial cell adherence and endocytosis

*C. albicans* adherence to and endocytosis of by oral epithelial cells was quantified by a differential fluorescence assay as described previously (Park et al. 2005). Briefly, OKF6/TERT-2 cells were grown to confluency on fibronectin-coated circular glass coverslips in 24-well tissue culture plates. They were infected with 2 × 10^5^ yeast-phase *C. albicans* cells per well and incubated for 2.5 h, after which they were washed, fixed, stained, and mounted inverted on microscope slides. The coverslips were viewed with an epifluorescence microscope, and the number of adherent and endocytosed organisms per high-power field (HPF) was determined, counting at least 100 organisms per coverslip. Each experiment was performed at least three times in triplicate. For MPO analysis during OPC, the tongue homogenates from immunocompromised mice a 3 d post-infection were clarified by centrifugation, and stored at −80°C. The MPO concentration was determined using a commercial ELISA kit (Hycult Biotech).

### Measurement of cytokines in immunocompromised and immunocompetent mice

#### *C. albicans* Als3 surface expression

Flow cytometry was used to analyze the surface expression of Als3 on C. *albicans* strains, using our previously described method (Phan et al. 2007; Fu et al. 2013). Briefly, after fixing C. *albicans* cells in 3% paraformaldehyde and blocking with 1% goat serum, the cells were incubated with a rabbit polyclonal antiserum raised against the recombinant N-terminal region of Als3. Next, the cells were rinsed and incubated with a goat anti-rabbit secondary antibody conjugated to Alexa Fluor 488 (Invitrogen). Cell sorting was performed with a FACSCaliber flow cytometer (Becton, Dickinson). Fluorescence data for 10,000 cells of each strain were collected.

#### Neutrophil killing

The susceptibility of the various C. *albicans* strains to neutrophil killing was determined as described previously (Solis et al. 2017). Briefly, neutrophils were straind from the blood of healthy volunteers and mixed with an equal number of *C. albicans* cells in RPMI 1640 medium plus 10% fetal bovine serum. After a 3 h incubation at 37°C in 5% CO_2_, the neutrophils were lysed by sonication, and the number of viable *C. albicans* cells was determined by quantitative culture. Each experiment was performed three times in triplicate (different donors).

#### Statistics

Data were compared by Mann-Whitney or unpaired Student’s t test using GraphPad Prism (v. 6) software. P values of < 0.05 were considered to be statistical significant.

#### Quantitative PCR (qPCR) to assess chromosome copy number

To check that the trisomies were maintained during inoculum preparation and throughout the course of infection we used qPCR for 4 markers along each of the 8 chromosomes to determine ploidy. The PCR primers are listed in Table S3 qPCR was performed on gDNA extracted from single colonies and from streaks of the inoculum and for strains recovered from the tongues of infected mice (Table S1). Streaks or single colonies were transferred from original plates directly into either 750 µl (streaks) or 150 µl (single colonies) of 50% glycerol, and 100 µl were used for gDNA extraction without additional culturing. Resulting gDNA amounts were sufficient for qPCR.

#### Whole genome sequencing and Sanger sequencing

Genomic DNA was isolated with phenol chloroform, as described previously (Selmecki et al. 2015). Libraries were prepared using the NexteraXT DNA Sample Preparation Kit following the manufacturer’s instructions (Illumina). Libraries were purified with AMPure XP beads (Agencourt) and library concentration was quantified using a Bioanalyzer High Sensitivity DNA Chip (Agilent Technologies) and a Qubit High Sensitivity dsDNA fluorometric quantification kit (Life Technologies). DNA Libraries were sequenced using paired end 2 × 250 flow cells on an Illumina MiSeq (Creighton University). Copy number and allele status was visualized using YMAP (Abbey et al. 2014). Fastq files were aligned using an in-house sequence analysis pipeline (Li and Durbin 2009; Li et al. 2009; Li and Durbin 2010) and custom Python scripts. The progenitor strain and any of the evolved (Bolger et al. 2014) strains were analyzed using Mutect (Cibulskis et al. 2013) resulting in individual output files containing *de novo* SNPs that were acquired by the evolved strains. SNP regions were validated by eye in IGV. For non-synonymous de novo SNPs, primer pairs were generated with Primer 3 (Untergasser et al. 2012) (Table S3) to yield PCR products of about 400 bp. SNPs were confirmed by Sanger sequencing of amplified products as described (Selmecki et al. 2006).

#### Spot dilution assays

Strains were streaked onto YPD plates and incubated for 3 days at 30°C. Single colonies were transferred to 3 ml YPD broth and grown overnight at 30°C in a roller incubator. Cells were spun down, washed twice with PBS buffer and resuspended in 1ml PBS. The number of cells/ml was counted using a hemacytometer and all strains were adjusted to 1 × 10^9^ CFU/ml. For each strain, five microliters of a five-fold dilution series were spotted onto YPD, YPD light (0.2% glucose), YPD supplemented with 1 M NaCl, 150 µM farnesol, 40 or 80 µg/ml Congo Red, 40 or 80 µg/ml Calcofluor White, 0.1 or 0.2% SDS, 10, 20, 40, or 100 mM mM H_2_O_2_, 15 ng/ml Rapamycin, Casitone media supplemented with 4 μg/ml fluconazole, or 0.1 μg/ml caspofungin, 3 mM Protamine sulfate, RPMI supplemented with 0.2% glucose and Spider medium (Azadmanesh et al. 2017). Four sets were prepared for each medium, and sets were incubated at room temperature 30°C, 37°C, and 42°C, respectively, monitored for growth and colony appearance and photographed on days 2, 3 and 6.

#### Extracellular lipase/phospholipase activity

To assay lipase and phospholipase hydrolytic activity, five microliters of the 1 × 10^9^ CFU/ml stock was spotted onto egg yolk agar plates (EYA; 10% egg yolk emulsion, 1% peptone, 1.5% agar, 3% glucose, 5.73% NaCl, 0.055% CaCl_2_) (Noumi et al. 2010). Spots were allowed to dry and plates were incubated at 37°C for up to 5 days, photographed on day 5 and images were analyzed with ImageJ. For each strain, twenty measurements of the extent of the clearing around each spot were taken. To determine whether the differences in clearing were significantly different from the parental strain and between strains unpaired Student’s t test were done using GraphPad Prism (v. 6) software. A p-value of < 0.05 was considered significant.

## Data availability

WGS data have been deposited at the SRA database under accession number SRP126179 (Temporary submission ID SUB3297543).

## Acknowledgements

AMD, GC, EL, and AF were funded by NIH grant R15 AI090633 to AF. This work was supported in part by NIH grants R01DE022600 and R01AI124566 to SGF., and grant K99DE026856 to MS. AMS was funded by Nebraska Tobacco Settlement Biomedical Research Development New Initiative Grant (LB692), NE-EPSCoR First Award, Nebraska Department of Health and Human Services (LB506-2017-55), CURAS Faculty Research Fund Award, and an NIH-NCRR COBRE grant P20RR018788 sub-award. JB was funded by an European Research Council Advanced Award 340087 (RAPLODAPT).

---

**Fig. S1.**
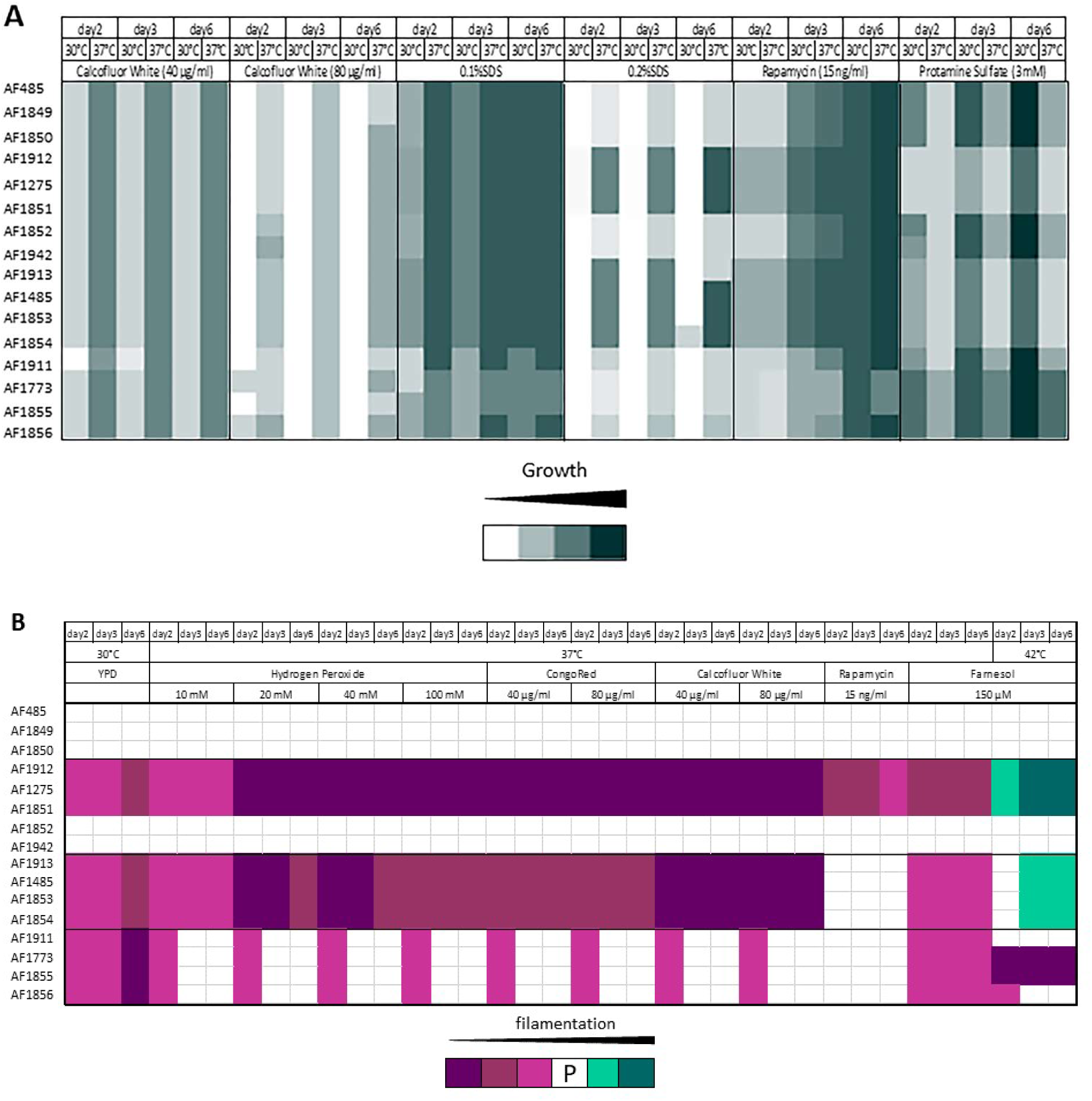
Summary of phenotypic evaluation of strain lineages. Shown are the conditions for which differences in growth (A) and spot morphology (B) were seen either between the progenitor and the trisomic strain(s) or between the different trisomic strains. Data is arranged by the day the plates were scored, the incubation temperature and growth medium. Strains and their derivatives are ordered as follows: progenitor, Chr6ABB, Chr6AAB, Chr5AAB. See Table S1 for strain information.

**Fig. S2.**
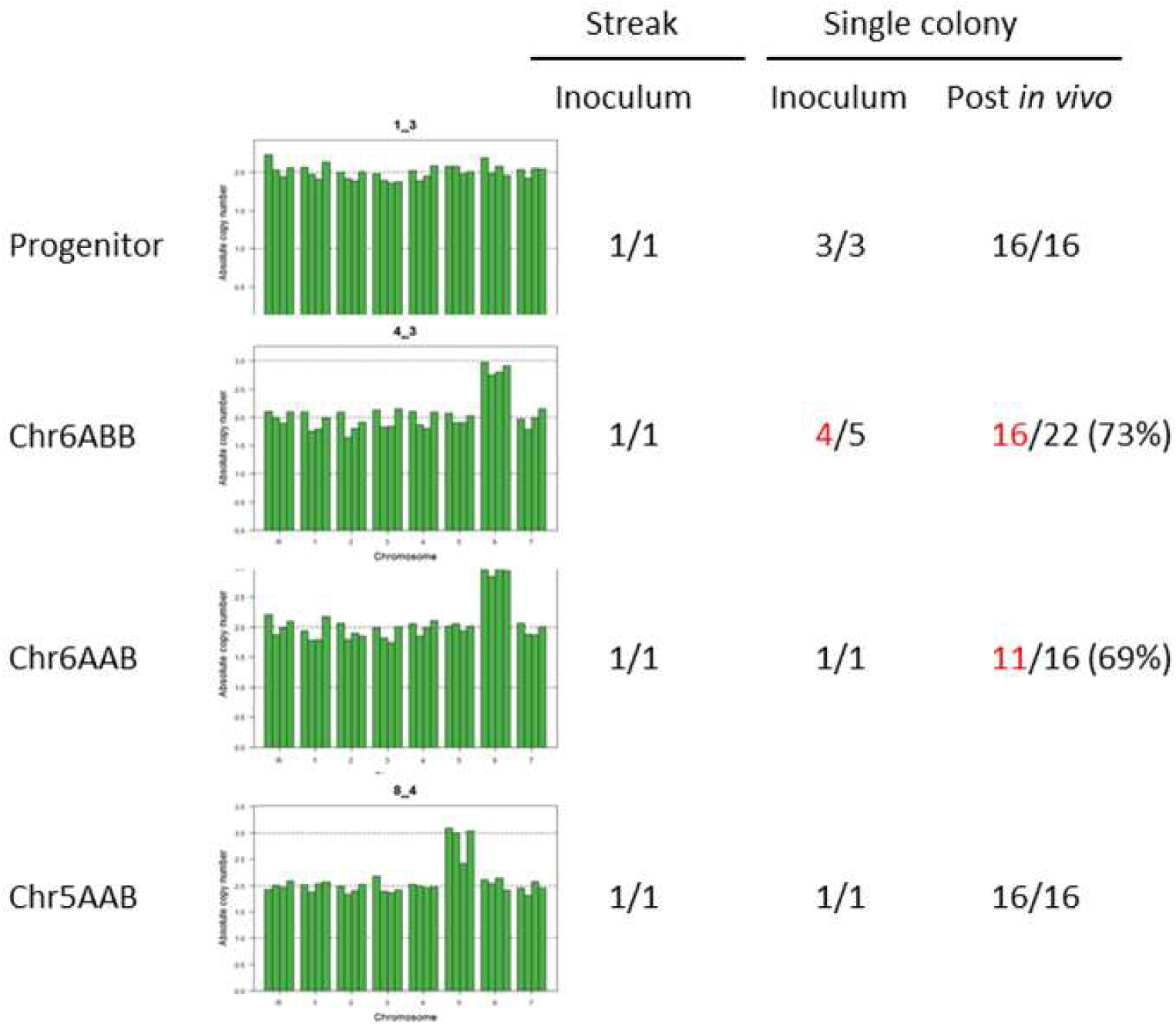
Variable retention of trisomies suggest population heterogeneity *in vivo.* Ploidy genotypes were stably maintained for the parent and the Chr5AAB lineage but not for either Chr6 lineage. The ploidy genotypes of the indicate strains was determined by qPCR for 4 markers along each of the 8 chromosomes.

