## Supplemental Table 1 for "Selection of *Candida albicans* Trisomy during Oropharyngeal Infection Results in a Commensal-Like Phenotype"

Table S1. Strains used in this study

| **Strain** | **Strain ID** | **parent** | **Genotype** | **Reference** |
| --- | --- | --- | --- | --- |
| AF485 | YJB9318 | RM1000 #2 | *gal1*::*URA3*/*GAL1,HIS1* | {Forche, 2018 #28629} |
| AF1911 | O3C10 | YJB9318 | Gal^+^, Chr5AAB | {Forche, 2018 #28629} |
| AF1912 | O4F7 | YJB9318 | Gal^+^, Chr6AAB | {Forche, 2018 #28629} |
| AF1913 | O4F8 | YJB9318 | Gal^+^, Chr6ABB | {Forche, 2018 #28629} |
| AF1275 | O4F7-BC23 | AF1912 | NEUT5L::NAT1-BC23 | This study |
| AF1485 | O4F8-BC29 | AF1913 | NEUT5L::NAT1-BC29 | This study |
| AF1773 | O3C10-BC3 | AF1911 | NEUT5L::NAT1-BC3 | This study |
| AF1849 | P1F1 | YJB9318 | see AF485 | This study |
| AF1850 | P1G2 | YJB9318 | see AF485 | This study |
| AF1851 | P2F8 | AF1275 | see AF1275 | This study |
| AF1852 | P2G1 | AF1275 | see AF1275 | This study |
| AF1942 | P2C1 | AF1275 | see AF1275 | This study |
| AF1853 | P3F7 | AF1485 | see AF1485 | This study |
| AF1854 | P3G7 | AF1485 | see AF1485 | This study |
| AF1855 | P4F8 | AF1773 | see AF1773 | This study |
| AF1856 | P4G7 | AF1773 | see AF1773 | This study |
