## Supplemental Table 3 for "Selection of *Candida albicans* Trisomy during Oropharyngeal Infection Results in a Commensal-Like Phenotype"

| Table S3. Primers used to confirm high confidence non-synonymous SNPs | | |
| --- | --- | --- |
| Primer ID | chromosome | primer sequence (5' to 3') |
| 792_orf19.7245_Chr1_3157737-F | 1 | CATGCACAGAACGAGATTGG |
| 793_orf19.7245_Chr1_3157737-R | 1 | ACGAAGATTGTGGGATCGAC |
| 786_VPS53_Chr1_20765-F | 1 | TGGTATCGCCATTGGTAAGC |
| 787_VPS53_Chr1_20765-R | 1 | CCGTGGGACATTTCTAATGC |
| 956_HIR1_Chr2_41910-F | 2 | CCTGACGCTAGTCTGGTTCC |
| 957_HIR1_Chr2_41910-R | 2 | GCAGTTCCATTGGACCAGTT |
| 611_orf19.4610_Chr4_359464_R | 4 | GGAGAGACAGGAGCAGGTTC |
| 612_orf19.4610_Chr4_359976_F | 4 | AGTGCCACCAAGACACACTT |
| 828_PMT1_Chr7_626471-F | 7 | CCTTGAGTATCAACGTGACGAA |
| 829_PMT1_Chr7_626471-R | 7 | CCATTCACCTTTGGCTGGTA |
| 832_RAD26_ChrR_1728166-F | R | TTGAATAGGCGTACCCGATAA |
| 833_RAD26_ChrR_1728166-R | R | CCAATGGGTCAATGAGTTCC |
| 609_HSP104_ChrR_1791692_F | R | TTGGCAAAACAACAGGCCAA |
| 610_HSP104_ChrR_1791100_R | R | CCAACACCAGCATCACCAATC |
