## Supplemental Figure 1 for "Selection of *Candida albicans* Trisomy during Oropharyngeal Infection Results in a Commensal-Like Phenotype"

Fig. S1. Summary of phenotypic evaluation of strain lineages. Shown are the conditions for which differences in growth (A) and spot morphology (B) were seen either between the progenitor and the trisomic strain(s) or between the different trisomic strains. Data is arranged by the day the plates were scored, the incubation temperature and growth medium. Strains and their derivatives are ordered as follows: progenitor, Chr6ABB, Chr6AAB, Chr5AAB. See Table S1 for strain information.

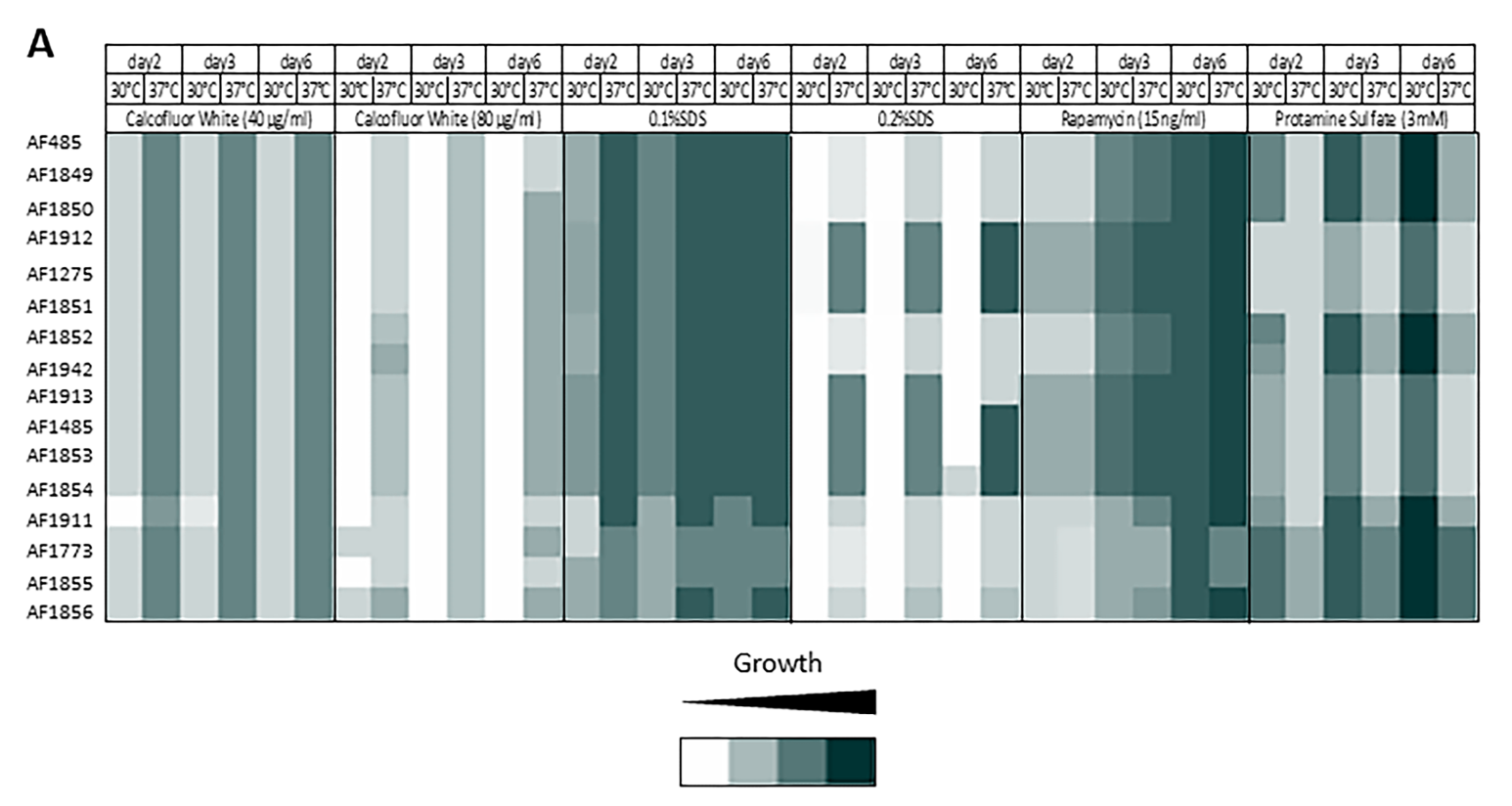

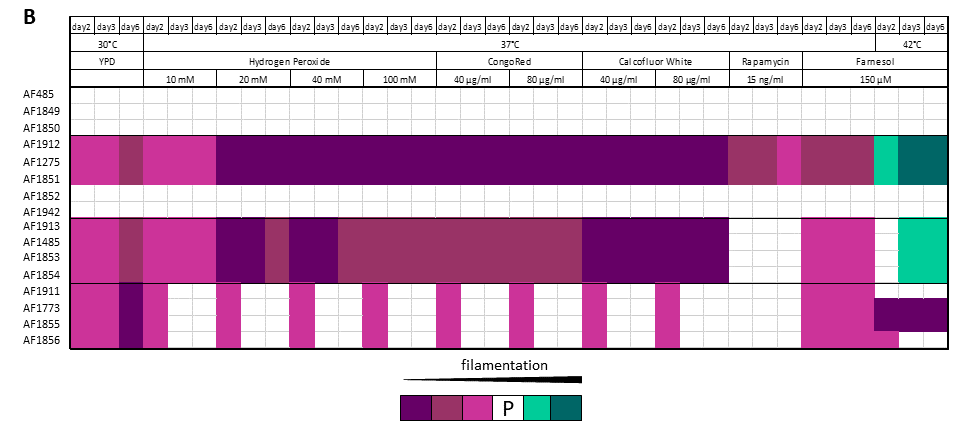
