## Supplemental Figure 2 for "Selection of *Candida albicans* Trisomy during Oropharyngeal Infection Results in a Commensal-Like Phenotype"

Fig.S2. Variable retention of trisomies suggest population heterogeneity *in vivo*. Ploidy genotypes were stably maintained for the parent and the Chr5AAB lineage but not for either Chr6 lineage. The ploidy genotypes of the indicate strains was determined by qPCR for 4 markers along each of the 8 chromosomes.

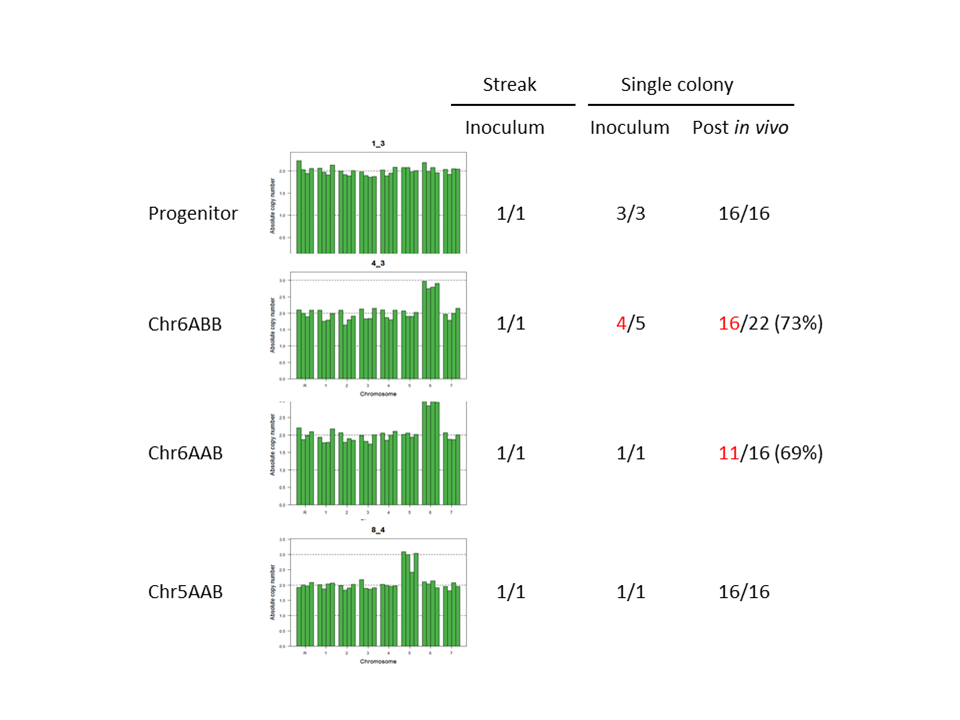
